# Single-nucleus RNA-seq identifies transcriptional heterogeneity in multinucleated skeletal myofibers

**DOI:** 10.1101/2020.04.14.041400

**Authors:** Michael J Petrany, Casey O Swoboda, Chengyi Sun, Kashish Chetal, Xiaoting Chen, Matthew T. Weirauch, Nathan Salomonis, Douglas P Millay

## Abstract

While the majority of cells contain a single nucleus, cell types such as trophoblasts, osteoclasts, and skeletal myofibers require multinucleation. One advantage of multinucleation can be the assignment of distinct functions to different nuclei, but comprehensive interrogation of transcriptional heterogeneity within multinucleated tissues has been challenging due to the presence of a shared cytoplasm. Here, we utilized single-nucleus RNA-sequencing (snRNA-seq) to determine the extent of transcriptional diversity within multinucleated skeletal myofibers. Nuclei from mouse skeletal muscle were profiled across the lifespan, which revealed the emergence of distinct myonuclear populations in postnatal development and their reactivation in aging muscle. Our datasets also provided a platform for discovery of novel genes associated with rare specialized regions of the muscle cell, including markers of the myotendinous junction and functionally validated factors expressed at the neuromuscular junction. These findings reveal that myonuclei within syncytial muscle fibers possess distinct transcriptional profiles that regulate muscle biology.

## Main

Skeletal muscle forms through the fusion of mononucleated muscle progenitor cells during development^1,2^. Mature, multinucleated myofibers are composed of different contractile and metabolic machinery that result in independent muscle fiber types (IIb, IIx, IIa and I)^3^. In addition to possessing fiber type diversity, skeletal muscle also contains a stem cell population (satellite cells) that can be activated upon a stimulus and ultimately fuse with myofibers or each other, thereby allowing regeneration^4^. Concepts to explain the requirement of multinucleation for skeletal muscle function include that each nucleus is capable of controlling a finite volume of cytoplasm and that regions of the myofiber perform specialized functions necessitating localized transcripts. Myofibers are known to exhibit functional specialization at postsynaptic endplates (neuromuscular junction: NMJ), where a coordinated transcriptional program is responsible for formation and maintenance of the synaptic apparatus^5-8^, although much remains unknown regarding the full network of factors responsible for NMJ development and function. Muscle-tendon connection sites (myotendinous junction: MTJ) are known to exhibit structural specialization^9^, but a transcriptionally unique population of myonuclei localized at the MTJ has not been defined. Whether additional regions of compartmentalization are present in skeletal muscle is also not understood, likely because of technical limitations related to assessing transcription in syncytial cells. Stochastic transcriptional pulsing of particular genes has been reported in myofibers, but it is unknown whether this phenomenon reflects true divergence of myonuclear identities^10^. Transcriptional diversity may also be present in physiological contexts that elicit myonuclear accretion from satellite cells, such as in exercise^11^. Overall, the full extent of myonuclear heterogeneity achieved within syncytial myofibers remains unknown but understanding this phenomenon will elucidate mechanisms that control muscle development and function.

We sought to gain insight into the range of myonuclear heterogeneity in mammals, through profiling of nuclei from mouse skeletal muscle using single-nucleus RNA-sequencing (snRNA-seq). Nuclei from 5-month mouse tibialis anterior muscle from wild-type mice were purified and the 10X Chromium system was used to build libraries for sequencing (Fig. 1a). We profiled 8331 nuclear transcriptomes, from which unbiased clustering revealed all major cell types expected in skeletal muscle, including myonuclei, satellite cells, fibroadipogenic progenitors (FAPs), endothelial cells, smooth muscle cells, tenocytes, and immune cells (Fig. 1b). Non-myonuclear clusters were identified through expression of canonical marker genes including *Pecam1* (endothelial cells), *Myh11* (smooth muscle), *Dcn* (FAPs), *Mkx* (tenocytes), and *Ptprc* (immune cells) (Extended Data Fig. 1, 2). We identified several clusters of myonuclei belonging to multinucleated muscle fibers, characterized by expression of numerous canonical myonuclear transcripts such as *Tnnt3, Mybpc1*, and *Mybpc2* (Extended Data Fig. 2), and constituting a population essentially excluded by conventional single-cell analyses of skeletal muscle^12,13^. Myonuclei could be further distinguished by muscle fiber type, determined by expression of distinct members of the myosin heavy chain gene family. The tibialis anterior is a fast-twitch muscle known to express predominantly the fast myosin heavy chain isoforms *Myh4* (Type IIb fibers) and *Myh1* (Type IIx fibers)^14^. Consistent with this, the four largest clusters of myonuclei were grouped by expression of these two markers (Fig. 1c). A small number of *Myh2*^+^ myonuclei (Type IIa) were also clustered upon higher-dimensionality analysis (Extended Data Fig. 3)^15^, although Type IIa myofibers are known to comprise only a small minority within the tibialis anterior^14^.

**Fig. 1.**
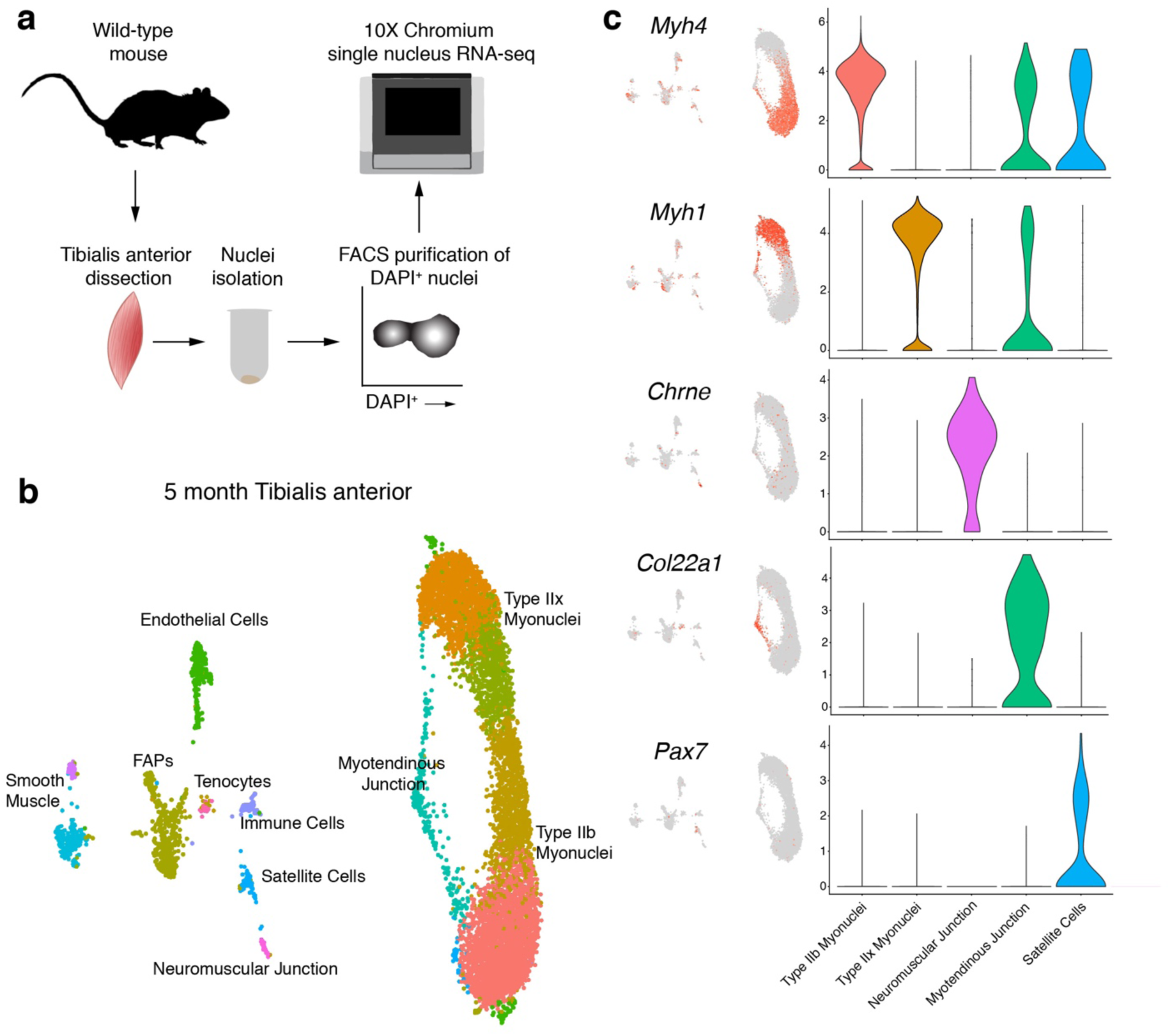
snRNA-seq of mouse tibialis anterior muscle at 5 months of age. **a**, Schematic for nuclei purification and sequencing from mouse skeletal muscle. **b**, Unbiased clustering of snRNA-seq data represented on a UMAP. **c**, UMAP and violin plots showing gene expression for myonuclear populations including *Myh4* (Type IIb myonuclei), *Myh1* (Type IIx myonuclei), *Chrne* (neuromuscular junction), *Col22a1* (myotendinous junction), and *Pax7* (satellite cells). Intermediate myonuclear clusters were combined with their respective fiber type myonuclei for generation of violin plots. The y-axis shows expression level as probability distribution across clusters.

Myonuclei that clustered outside of fiber type included those of the neuromuscular and myotendinous junctions. 0.8% of myonuclei were identified as belonging to the neuromuscular junction (NMJ), where postsynaptic transcriptional specialization is known to occur. NMJ nuclei were positive for known canonical markers such as *Chrne, Etv5, Colq*, and *Musk* (Fig. 1c and Extended Data Fig. 4a), but also expressed numerous genes not previously associated with the NMJ, with a total of 345 significantly upregulated genes. Myotendinous junction (MTJ) nuclei comprised 3.6% of myonuclei and were notable for expression of *Col22a1* and *Ankrd1* (Fig. 1c and Extended Data Fig. 4b), whose expression has been reported to concentrate at the ends of muscle fibers^16,17^. Despite data showing that myonuclei may cluster near the MTJ^18,19^, robust evidence of a unique nuclear population with a specific transcriptional program has not been shown. Here, we identified 291 uniquely upregulated genes, and genes previously not associated with the MTJ included *Slc24a2* and *Adamts20* (Extended Data Fig. 4b). The source of *Col22a1* at the ends of myofibers is not clear, but recent evidence indicates contribution from a tenocyte population^20^. Our data show *Col22a1* and myosins are co-expressed, as well as *Col22a1* expression in tenocytes (Fig. 1c), suggesting both myonuclei and tenocytes contribute this extracellular matrix protein. These data indicate strong spatial transcriptional heterogeneity of NMJ and MTJ myonuclei, suggesting that location within the syncytia and associated proximal signaling are major determinants of transcriptional profiles. In addition to transcriptional specialization at the NMJ and MTJ, we asked whether any further heterogeneity within fiber-type specific myonuclei was discernible. Sub-clustering of exclusively *Myh4*+ (Type IIb) or *Myh1*+ (Type IIx) myonuclei revealed multiple nuclear compartments with divergent transcriptional states (Extended Data Fig. 5a, b), raising the possibility of additional myonuclear sub-types with distinct functions.

To test if myonuclear heterogeneity is dependent upon particular muscle group, we profiled 5389 nuclei from the slow-twitch soleus muscle. Here, the populations detected were consistent with those in the tibialis anterior but also included Type I myonuclei (*Myh7*^+^), which is characteristic of the soleus muscle (Extended Data Fig. 6a, b)^14^. Integration of the tibialis anterior and soleus data resulted in a comprehensive atlas of fiber-type expression profiles and showed that Type I myonuclei are the most divergent likely owing to unique activities of this muscle group (Extended Data Fig. 6c, d).

While our analysis of 5 month old muscle suggests the possibility of previously unknown myonuclear diversity, outside of the NMJ and MTJ, it was not clear if the level of diversity indicated true functional compartmentalization. We asked whether more prominent myonuclear heterogeneity might be evident during postnatal development. Robust accrual of myonuclei during postnatal development typically ceases by 21 days in healthy mice, by which time the myogenic differentiation program has been switched off and myofibers possess nearly their full complement of myonuclei^21^. However, muscle growth and maturation continue beyond this point until mice reach adulthood, and how this process of muscle growth impacts transcription in accrued myonuclei is unclear. We profiled 11,552 nuclear transcriptomes from post-natal (P) day 21 tibialis anterior muscle to assess transcriptional dynamics of maturing myonuclei. P21 myonuclei clustered in a pattern distinct from that of adult mice, and clusters were not solely reducible to fiber-type identity (Fig. 2a). In addition to Type IIb, Type IIx, NMJ, and MTJ myonuclei, two additional myonuclear populations were identified. One of the developmental clusters of myonuclei were characterized by expression of *Enah* and the other expression of *Nos1* (Extended Data Fig. 7a). Further clustering of only myonuclei showed more specific subpopulations that were clearly distinguished from the Type IIb and Type IIx myonuclei (Fig. 2b). Differential gene expression analysis revealed two clusters of *Ttn*^+^ myonuclei that less strongly expressed markers of end-state myonuclear differentiation (*Myh1, Myh4, Ckm)*, which we named transient states A and B (Fig. 2b, Extended Data Fig. 7b). These *Myh4*^*-*^/*Myh1*^*-*^ clusters showed a skeletal muscle-specific gene signature (Extended Data Fig. 8), but did not highly express myogenic markers (*Myod1, Myog*, and *Mymk*) suggesting they represent nuclei within myofibers (Fig. 2c). Three additional populations were identified: two displayed scattered expression of *Myh4* or *Myh1* (Fig. 2c and Extended Data Fig. 7b), which we assigned fiber type-like designations; and a small unknown *Meg3*^+^ population. Transient states A and B displayed unique transcriptional programs and were enriched for factors associated with early myofibrillogenesis, including *Nrap, Fhod3, Enah*, and *Flnc*, as well as the non-muscle myosins *Myh9* and *Myh10*, which have been postulated as components of “pre-myofibrils” laid down as scaffolds before mature muscle-specific myosins are assembled (Fig. 2d)^22-27^. Multiple transcription factors were enriched in transient myonuclei, including both known drivers of skeletal muscle differentiation (*Ifrd1, Nfat5, Mef2a*)^28-30^ and numerous TFs with no previous associations in muscle (*Ell, Creb5, Zfp697*) (Extended Data Fig. 7c). We performed TF binding site enrichment analysis on the upregulated genes within each cluster, and discovered that Atf3 motifs were significantly enriched near transcriptional start sites of transient state B marker genes (Extended Data Fig. 9a), and that *Atf3* transcripts were highly increased in that same population in our snRNA-seq data (Extended Data Fig. 9b). This provides evidence for a coordinated transcriptional regulatory program and suggests a previously unknown role for Atf3 in myonuclear function during development. Thus, these data show temporal myonuclear heterogeneity within skeletal muscle at P21 and suggests that the myonuclear transient states represent a distinctive, specialized stage in which coordinated transcriptional activity supports increased myofibrillogenesis.

**Fig. 2.**
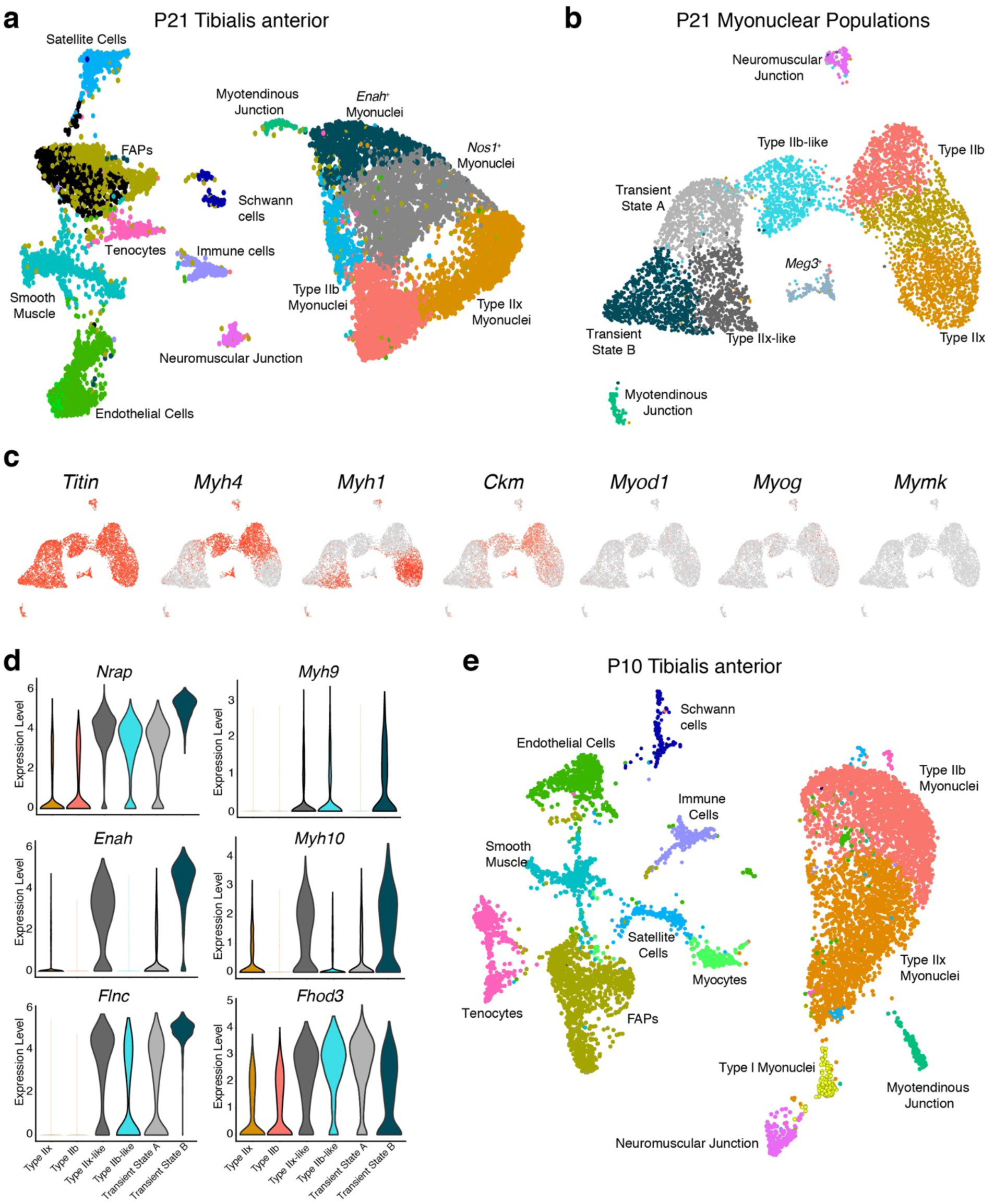
Temporal myonuclear heterogeneity revealed in developing muscle. **a**, Unbiased clustering of nuclei from postnatal (P) day 21 presented in a UMAP revealed all major populations in addition to unique myonuclei outside the canonical Type IIb and Type IIx myonuclei. These nuclei populations were marked by *Nos1* and *Enah* respectively. **b**, Sub-clustering of myonuclear populations from P21 muscle revealed transient myonuclear states. **c**, Feature plots assessing expression of mature muscle markers (*Ttn, Myh1, Myh4*, and *Ckm*) and differentiation markers (*Myod1, Myog, Mymk*) from the myonuclear populations in (b). **d**, Violin plots showing the transient myonuclei are enriched for a transcriptional profile associated with early myofibrillogenesis (expression of *Nrap, Enah, Flnc, Myh9, Myh10, Fhod3*). **e**, UMAP representing snRNA-seq data from a P10 tibialis anterior shows the presence of an activated muscle progenitor population (myocytes) but the absence of transient myonuclear states.

One interpretation of P21 transient myonuclear states is that they are the most recently fused nuclei and have not yet established their fiber type. In this scenario, these transient myonuclei could be in a more immature state and suggest that they progress through a post-fusion maturation program to establish their adult identity. Another possibility is that the transient myonuclear state is entrained after the majority of myonuclear accretion has occurred (around P21). To distinguish these possibilities, we generated 8611 nuclear transcriptomes from the tibialis anterior at P10 when fusion is ongoing and recently fused nuclei would be present. Here we were unable to detect the transient myonuclear populations and instead detected myonuclear populations similar to adult muscle (Fig. 2e). Interestingly, we did detect a new cluster of cells at P10, which were close to satellite cells, but were not present at P21 or 5 months of age. We hypothesized that this population represented myocytes (activated satellite cells) since fusion is ongoing at P10. We sub-clustered the satellite cell and myocyte populations and detected the presence of differentiation and fusion genes including *Myog* and *Mymk* in the myocyte population, while *Pax7* was enriched in the satellite cell population (Extended Data Fig. 7d). The presence of myocytes demonstrated ongoing fusion at P10 and the absence of this population at P21 confirmed that the majority of fusion was terminated in these samples. The lack of transient myonuclear populations at P10 indicates that they do not represent maturation immediately proximal to fusion, but instead appear in response to a physiological stimulus during the post-fusion growth of skeletal muscle.

In addition to functional diversity found in normal development, transcriptional heterogeneity can arise from pathological or compensatory processes associated with aging^31-33^. Skeletal muscle undergoes dramatic age-related changes with respect to functional deterioration and regenerative decline^34-37^, and so we next asked if the aging process impacted myonuclear transcription. We profiled 18,087 nuclei from 24-month and 30-month old tibialis anterior muscles (Fig. 3a, b). In 30-month muscle only, we observed emerging myonuclear populations that were not present in 5 or 24 months, including clusters expressing *Nr4a3, Ampd3*, and *Enah*, respectively (Fig. 3b). We integrated myonuclei from 5, 24, and 30 months and found a consistent aging-related gene expression signature including upregulation of *Nr4a3* and *Smad3* at both aged time-points (Fig. 3c), indicating that this may be a progressive feature of aging muscle. *Smad3* has been implicated during the muscle aging process^38,39^, while *Nr4a3* has a role for metabolic adaptations to exercise but has not been studied during aging^40,41^. smFISH for *Nr4a3* in the gastrocnemius muscle confirmed elevated expression in myonuclei in aged muscle, following a heterogeneous pattern (Fig. 3d). The *Ampd3*^+^ population were identified as myonuclei based on the presence of the myogenic regulatory factor *Myf6*, the pro-atrophy gene *Fbxo32*, and neuromuscular markers *Musk, Chrnb1, and Hdac4* (Extended Data Fig. 10). The cluster was also enriched for genes associated with the immune response, apoptosis, and proteasomal degradation (*Tnfrsf23, Traf3*, and *Psma5*) (Extended Data Fig. 10). Taken together, these data suggest that the *Ampd3*^+^ population could represent a denervated state that is dysfunctional (Extended Data Fig. 10). Surprisingly, the *Enah*^+^ myonuclear cluster was found to be highly similar to the *Enah*^+^ transient state B from P21 muscle, with numerous marker genes in common including *Atf3, Flnc*, and *Nrap* (Fig. 3e). This indicates that aging muscle unexpectedly can result in reactivation of a coordinated developmental transcriptional program, which may function as an attempt to compensate for age-related muscle decline.

**Fig. 3.**
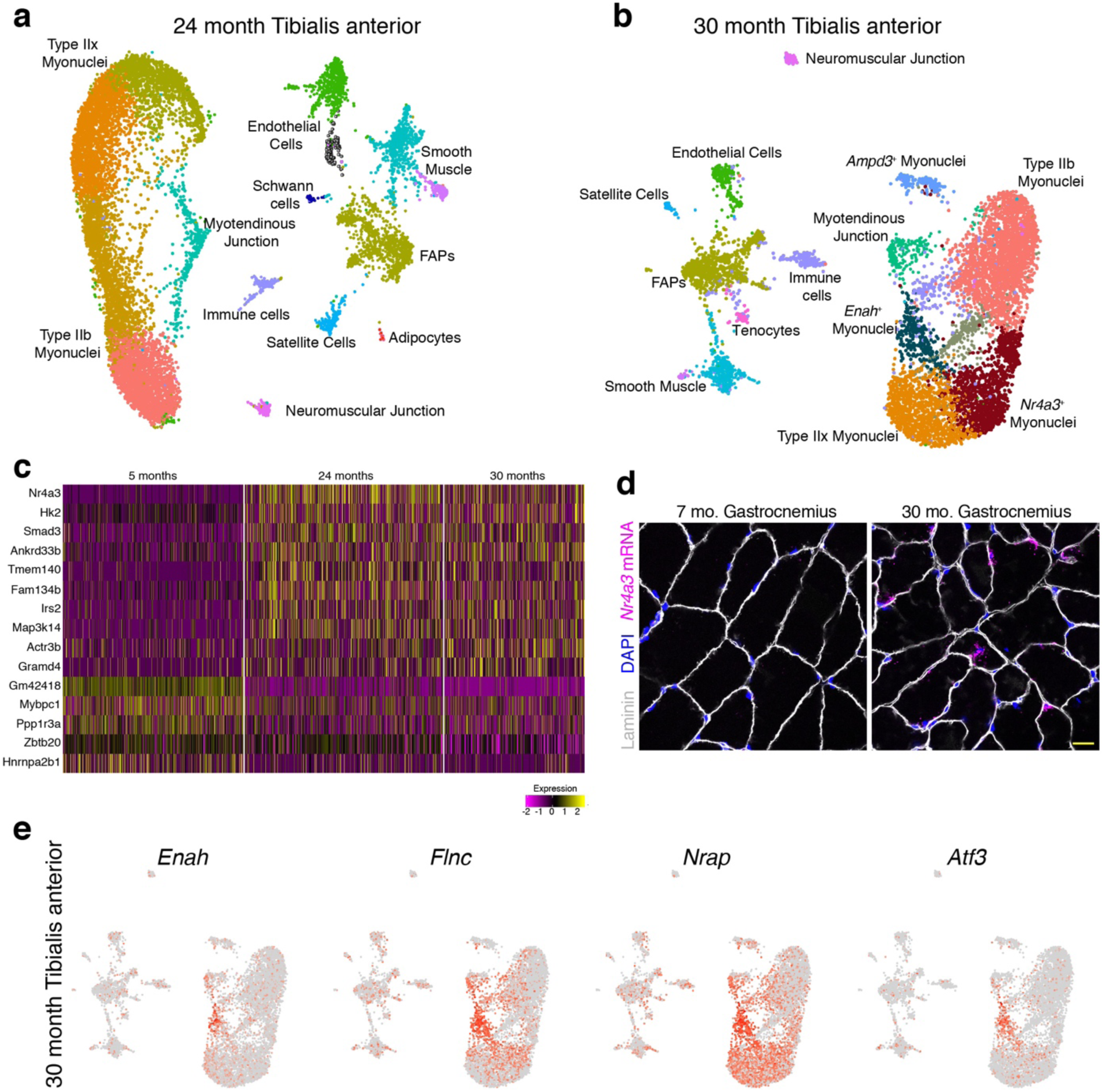
Uniform gene expression between myonuclei is disrupted in aged muscle. **a**, UMAP of snRNA-seq data from the tibialis anterior of a 24-month-old mouse displaying similar clusters as at 5 months of age. **b**, UMAP of snRNA-seq data from 30-month old muscle displaying the presence of unique clusters of myonuclei marked by *Ampd3, Enah*, and *Nr4a3*. **c**, Heatmap of integrated myonuclei from 5, 24, and 30-month tibialis anterior muscle showing a consistent transcriptional signature of aging muscle. Columns are individual nuclei belonging to each to time-point. **d**, Representative image of smFISH for *Nr4a3* validated its upregulation in myonuclei in 30-month skeletal muscle (n=3 for each group). Scale bar: 10 μm. **e**, Feature plots of key marker genes of the *Enah*^+^ population within 30-month myonuclei.

In addition to revealing the emergence of shared transcriptional responses at distinct stages of the life cycle, our data also establish an atlas of myonuclear transcriptomics and reveal transcripts associated with rare myonuclear populations such as the NMJ. To this end, an interactive data portal is publicly available at https://research.cchmc.org/myoatlas. Previous studies have elegantly uncovered unique gene expression in NMJ-associated nuclei which allowed for identification of fundamental molecular components of postsynaptic function ^42-44^. However, our data reveals numerous previously unreported genes associated with the NMJ, likely because our approach was able to resolve global transcription at the nuclear level. This represents a unique opportunity to further reveal mechanisms of postsynaptic development and function. To test the validity of newly-discovered NMJ marker genes, we confirmed localization of 4 of the genes not previously associated with the NMJ (*Ufsp1, Lrfn5, Ano4*, and *Vav3*) by smRNA-FISH and co-labeling of the NMJ marker acetylcholine receptor (AchR) (Fig. 4a). Moreover, we found that multiple top NMJ genes identified here showed significant changes in mRNA levels following hindlimb denervation, an expression pattern associated with postsynaptic regulation (Fig. 4b)^45^. While the expression of the majority of those genes increased after denervation, similar to the canonical NMJ factor *Musk*^*46*^, *Ufsp1* and *D430041D05Rik* exhibited a decrease in expression, while *Pdzrn4* did not change (Fig. 4b). To assess the functional relevance of these factors for NMJ formation and maintenance, we used an *in vitro* culture system in which C2C12 myoblasts, when plated on laminin, form aneural NMJ-like clusters that are marked by AChR (Fig. 4c)^47-49^. We treated C2C12 cells with a control siRNA or siRNAs targeting *Musk, Ufsp1, Vav3, B4galnt3*, or *Gramd1b*, then assayed for AChR clustering. Reduced expression of the target genes was observed after treatment with the appropriate siRNA (Fig. 4d). *Musk* reduction results in a blockade of AChR clustering (Fig. 4e), showing fidelity of the system^47^. Obvious AChR clustering abnormalities were not observed for cultures where *Vav3* or *B4galnt3* was reduced, but *Ufsp1* and *Gramd1b* were identified as regulators of the NMJ (Fig. 4e). Loss of *Gramd1b* resulted in reduced AChR clusters, whereas reduction of *Ufsp1* elicited an increase (Fig. 4f). These data indicate that *Ufsp1* is a negative regulator of AChR clustering and NMJ formation, consistent with its down-regulation following denervation. Reduction of *Ufsp1* or *Gramd1b* did not impact overall myogenesis or transcript levels of canonical NMJ genes (Fig. 4g), suggesting they specifically regulate the NMJ in a transcriptionally-independent manner. These data highlight the utility of nuclear-level resolution for the discovery of factors that regulate the biology of skeletal muscle.

**Fig. 4.**
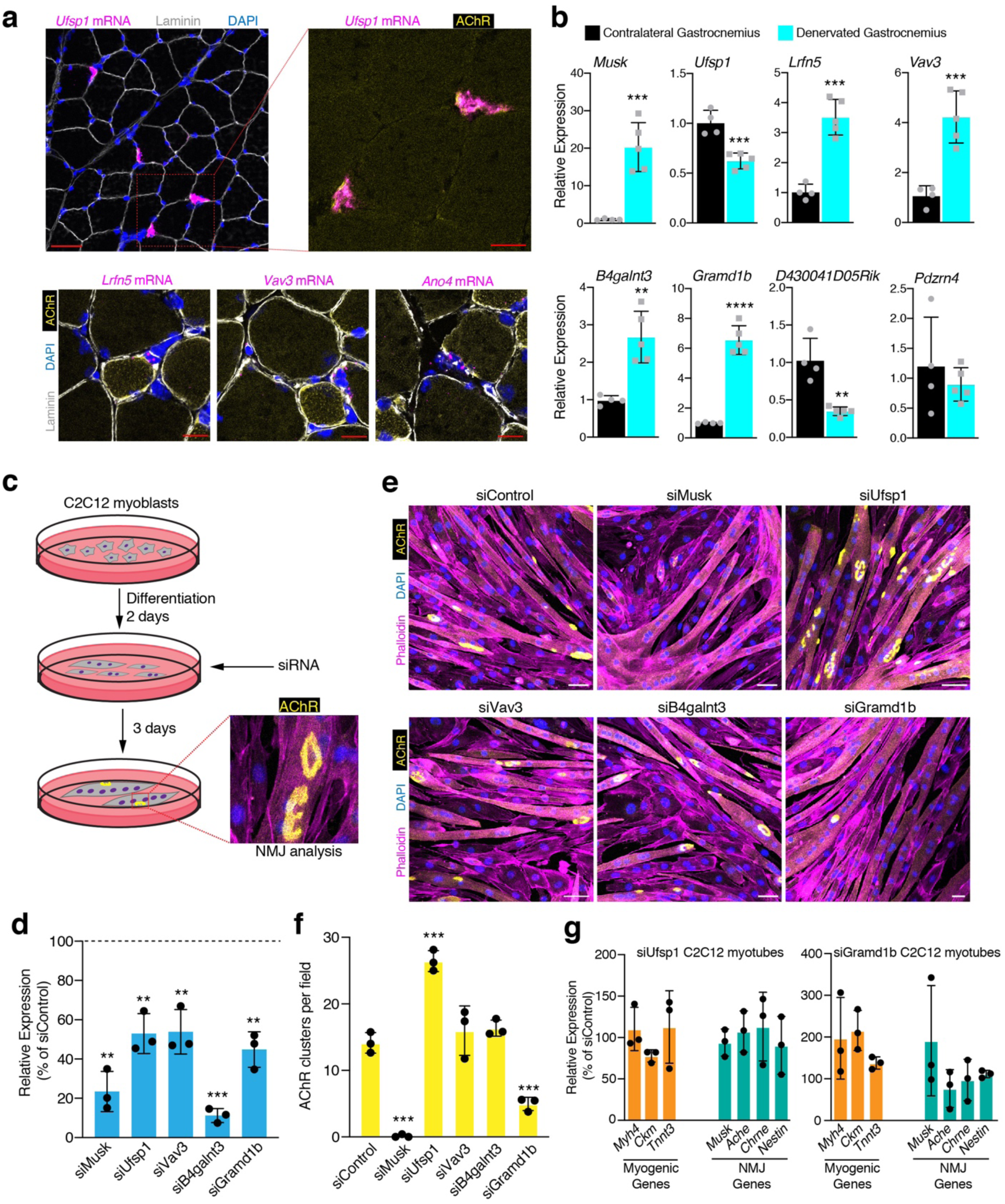
Discovery and functional characterization of novel NMJ genes. **a**, Representative images from smFISH for *Ufsp1, Lfrn5, Vav3*, and *Ano4* on skeletal muscle sections showing co-localization with the canonical NMJ protein, acetylcholine receptor (AChR) (n=3). AChR was visualized through α-bungarotoxin labeling. **b**, Quantitative real-time PCR (qPCR) analysis for the indicated genes not previously associated with the NMJ from normal and denervated muscle. **c**, Schematic for a siRNA screen in C2C12 myoblasts designed to test the function of novel NMJ genes. siRNA was transfected two days after differentiation and three days later α-bungarotoxin was used to analyze AChR clustering as a surrogate for NMJ formation. **d**, qPCR analysis for the genes targeted with siRNA. A scrambled siRNA was used as a control. **e**, Representative images of C2C12 myotube cultures after treatment with various siRNAs and staining with α-bungarotoxin. Cells were also stained with phalloidin and DAPI. **f**, Quantification of AChR clusters per field of view from (E). **g**, qPCR analysis for genes associated with myogenesis and NMJ formation. Scale bars: (a) 50 μm (top left panel), 10 μm (top right and bottom panels). Data in (b), (d), (f), and (g) are represented as mean ± standard deviation from three independent experiments. An unpaired t-test was used to determine statistical significance, ** P<0.01, ***P<0.001, ****P<0.0001.

Altogether, our single-nucleus profiling of skeletal muscle reveals intricate transcriptional dynamics of myonuclear states. An unexpected finding was the presence of unique transient myonuclear states in developing muscle after postnatal fusion and myonuclear accrual was terminated, and the reemergence of a shared transcriptional signature in aged muscle. We note that such transient states are likely real as combined analysis of all captures, using an independent approach and without batch correction, found overlapping cells in all clusters, indicative of temporally related cell states (Extended Data Fig. 11). We speculate that the transient state is determined by stage-specific growth signals following the tapering of developmental fusion, and is reactivated in aging muscle as a compensatory response to atrophy and diminished function. Our analyses also provide a comprehensive resource for further study of rare but essential myonuclear populations. We identify numerous novel genes for the specialized NMJ and MTJ compartments, and provide evidence that newly-discovered NMJ factors are functional regulators of postsynaptic function. While we analyzed all nuclei from muscle, approaches to enrich specifically for myonuclei (Kim M., Franke V., Brandt B., Spuler S., Akalin A., Birchmeier C. Single-nucleus transcriptomics reveals functional compartmentalization in syncytial skeletal muscle cells. Submitted to BioRxiv.) or to improve transcript detection levels may reveal further insights into myonuclear heterogeneity. Further experiments will be required to identify the key regulatory factors that drive the transient myonuclear state during development and aging, and the mechanisms that lead to assignment of certain nuclei to specialized compartments within a shared myofiber cytoplasm.

**Extended Data Fig. 1.**
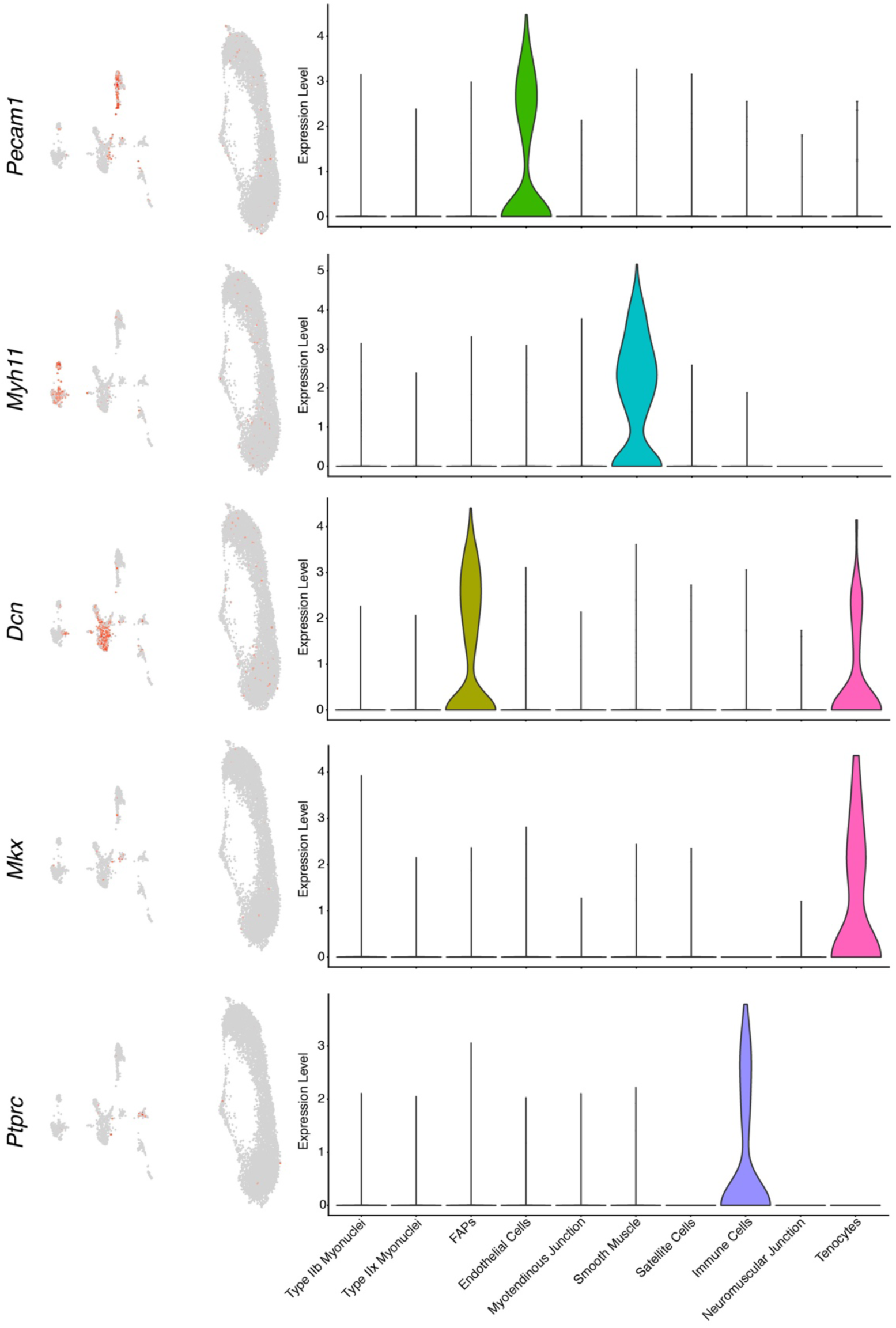
Identification of non-muscle nuclei in 5-month tibialis anterior snRNA-seq dataset. Feature and violin plots for markers of endothelial cells (*Pecam1*), smooth muscle (*Myh11*), fibroadipogenic progenitors (FAPs) (*Dcn*), tenocytes (*Mkx*), and immune cells (*Ptprc*). Type IIb and Type IIx myonuclei were combined with their intermediate clusters for generation of violin plots.

**Extended Data Fig. 2.**
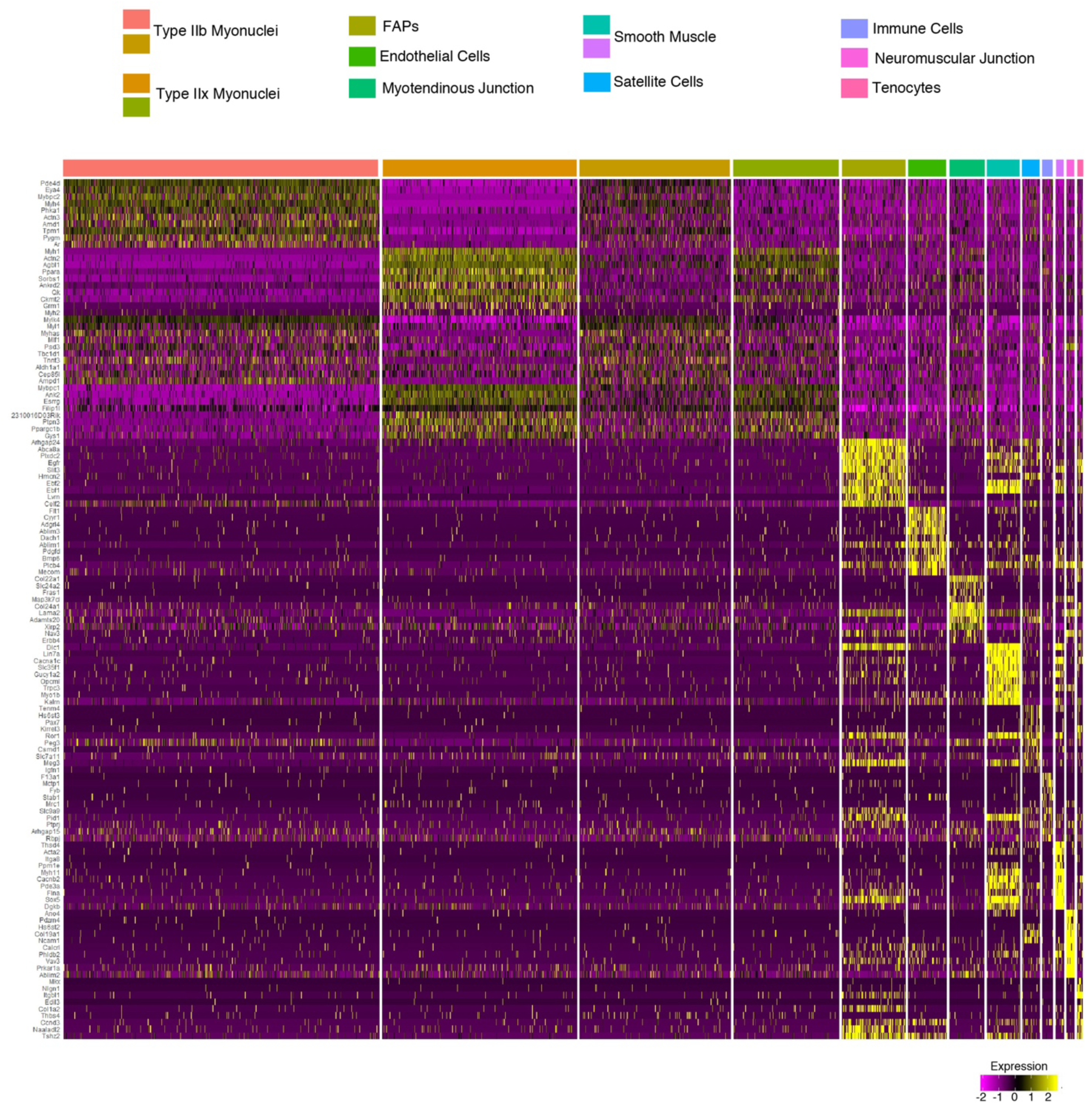
Top differentially expressed marker genes for clusters in 5-month tibialis anterior muscle. Normalized expression (Z-score) heatmap showing the top differentially expressed genes among the 13 clusters identified by snRNA-seq. Columns are individual nuclei belonging to each of the clusters labeled above.

**Extended Data Fig. 3.**
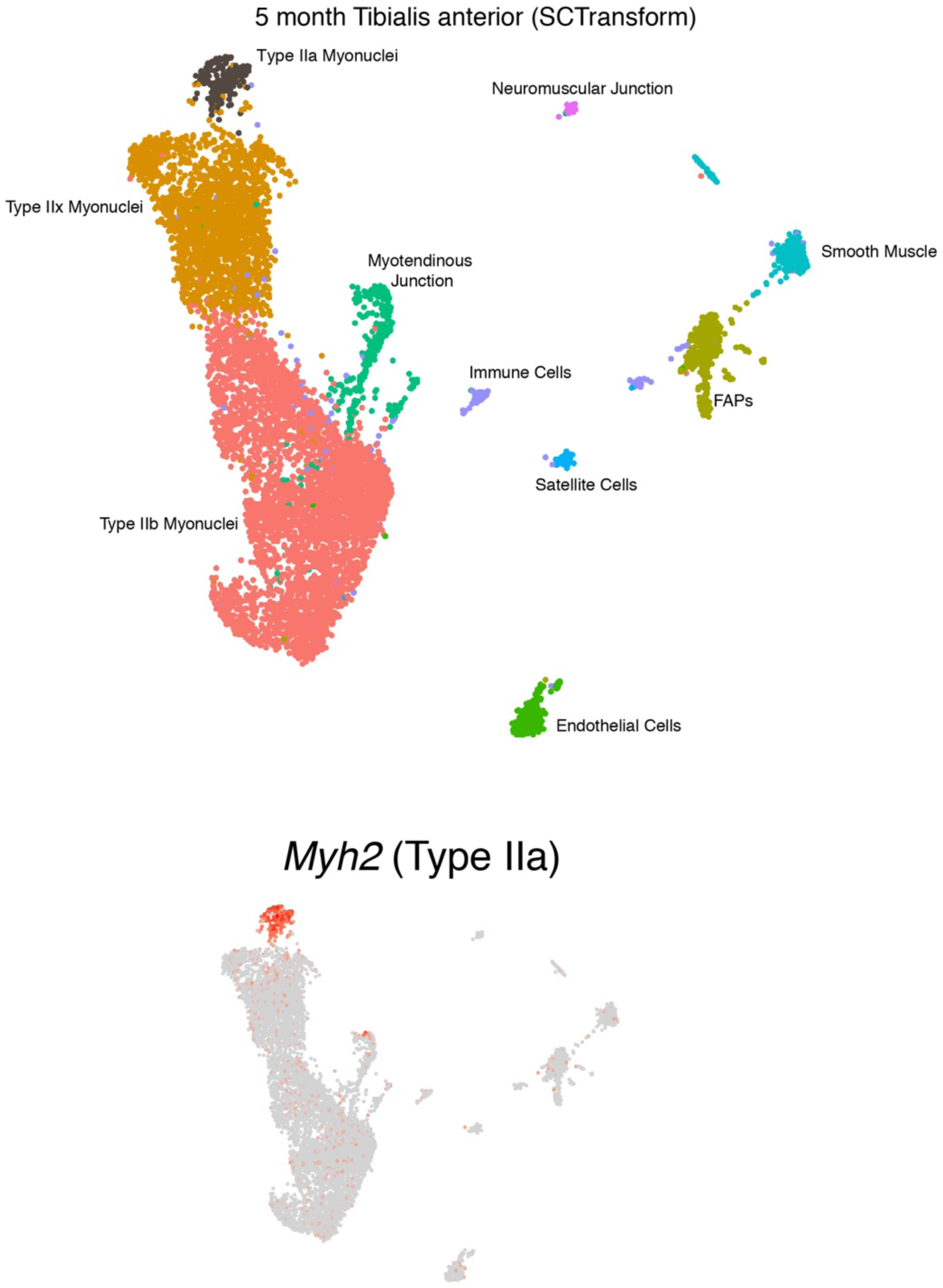
Higher dimensionality clustering of nuclear sequencing data through use of the SCTransform function. This analysis revealed the presence of myonuclei positive for *Myh2* (Type IIa). Type IIa fibers are known to comprise a small percentage of the fiber types in the tibialis anterior.

**Extended Data Fig. 4.**
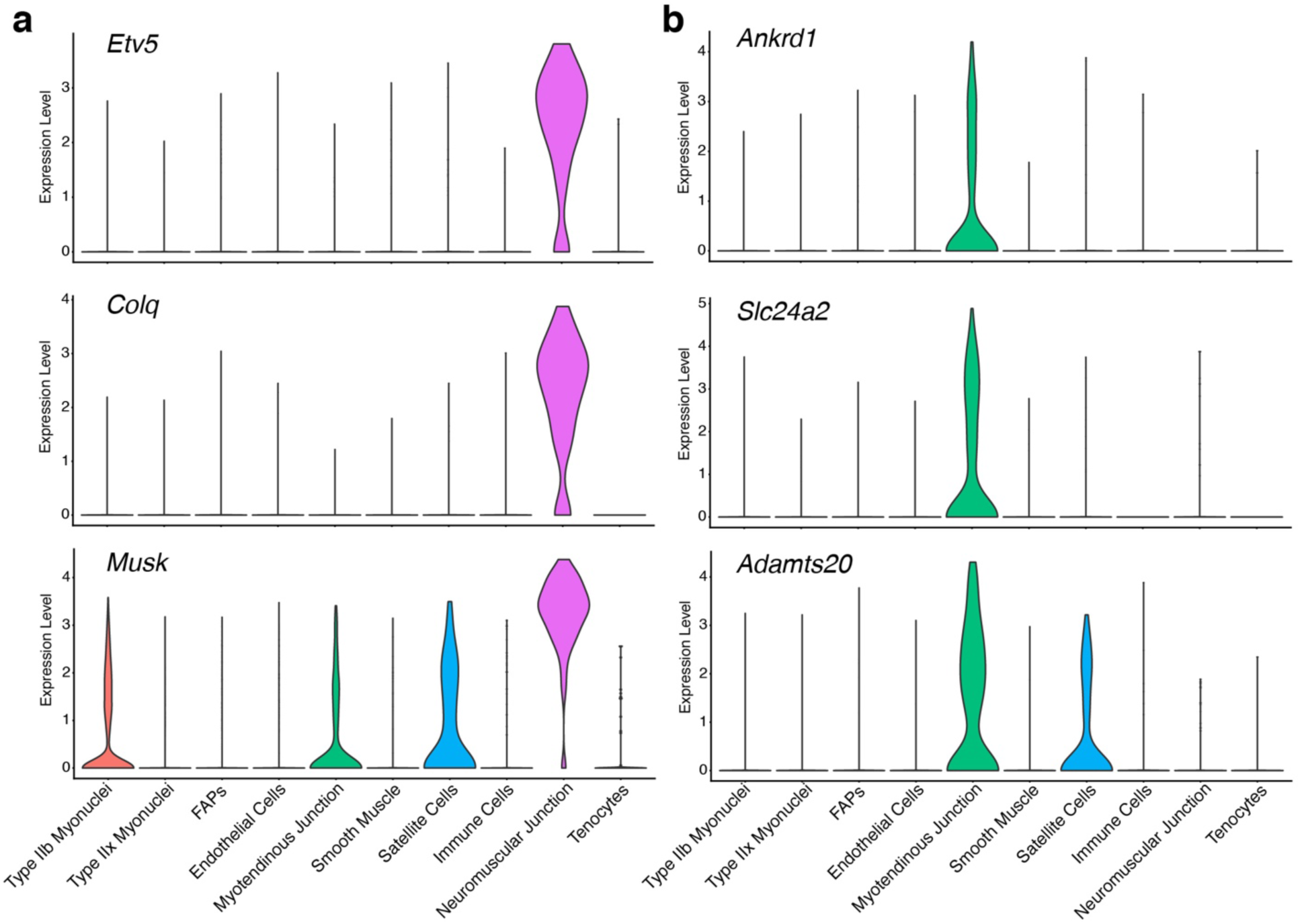
Identification of known genes and genes previously not associated with the neuromuscular junction (NMJ) and myotendinous junction (MTJ). **a**, Violin plots for the NMJ-enriched genes including *Etv5, Colq*, and *Musk*. **b**, Violin plots for *Ankrd1, Slc24a2*, and *Adamts20*, which we identified as being enriched in MTJ myonuclei.

**Extended Data Fig. 5.**
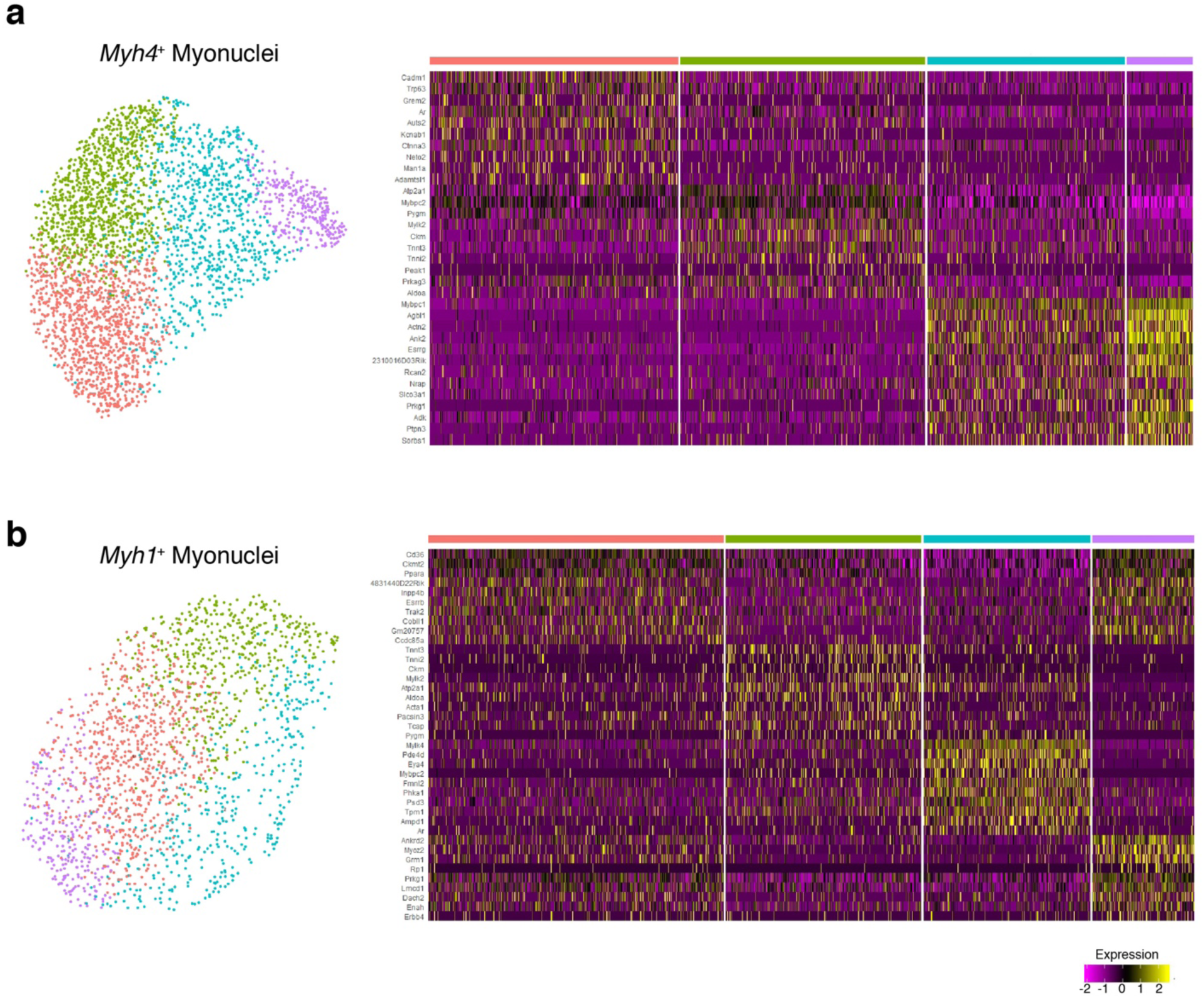
Fiber type-specific myonuclei show diversity of transcriptional states. **a**, Sub-clustering with UMAP projection and normalized expression (Z-score) heatmap generation of *Myh4*^+^/*Myh1*^-^/*Myh2*^-^ (exclusively Type IIb) myonuclei. **b**, UMAP and heatmap of *Myh1*^+^/*Myh4*^-^/*Myh2*^-^ (exclusively Type IIx) myonuclei. Both myonuclear types show sub-clusters divergent gene expression states. Columns are individual nuclei.

**Extended Data Fig. 6.**
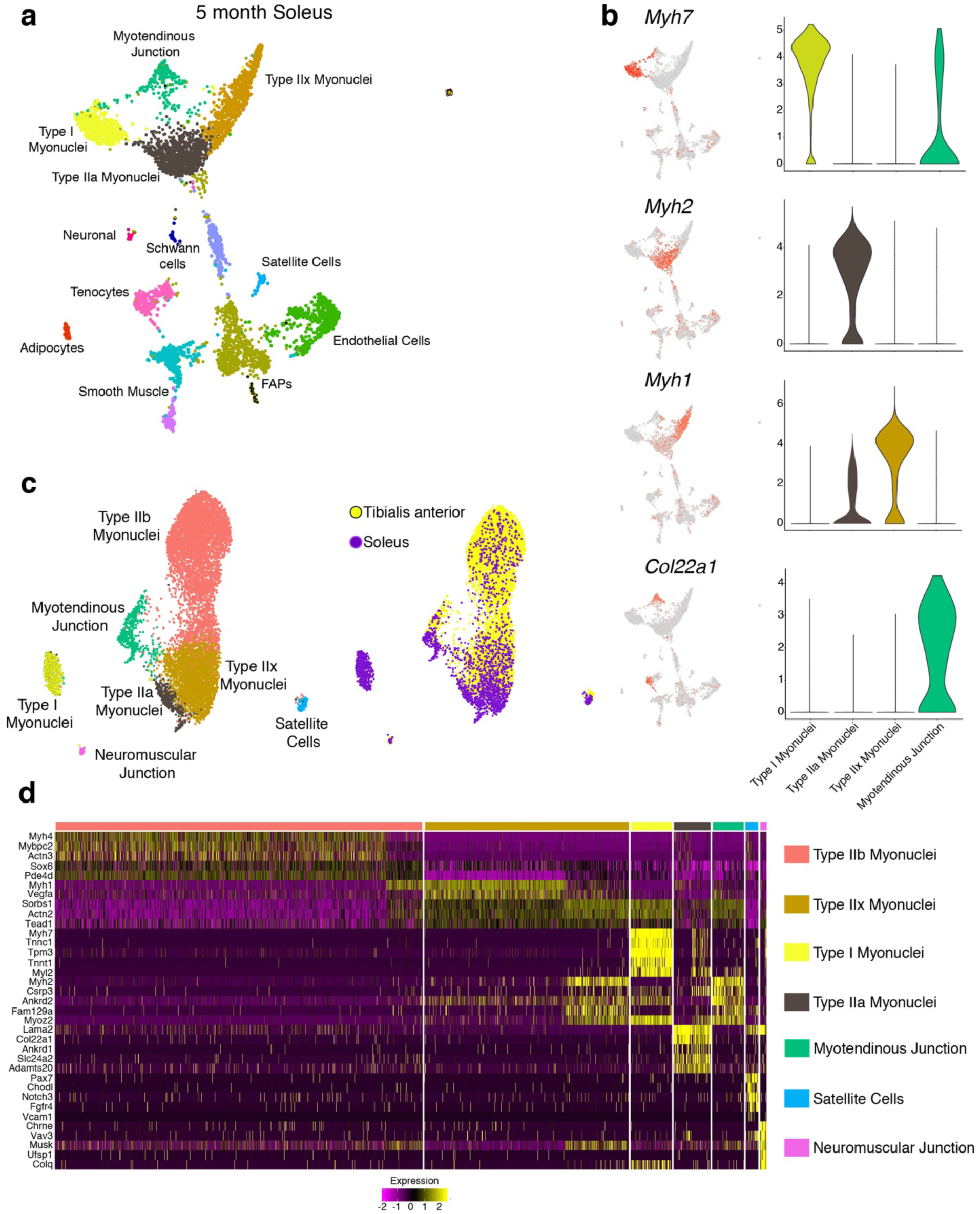
Myonuclei within all fiber types exhibit uniform transcription. **a**, Unbiased clustering of nuclei represented in a UMAP from the soleus muscle showing Type I, Type IIa, and Type IIx myonuclei. **b**, Feature and violin plots for *Myh7, Myh2*, and *Myh1* showing enriched expression in the expected fiber types. *Col22a1* also marks the myotendinous junction in slow-twitch muscle. **c**, Integration of myonuclear populations from the tibialis anterior and soleus muscles showing that Type I myonuclei are the most divergent myonuclear population. The right panel shows if the nuclei in the left panel originates from the tibialis anterior or soleus. **d**, Heatmap showing the top genes expressed in the muscle-related nuclear populations present in the integrated tibialis anterior/soleus dataset.

**Extended Data Fig. 7.**
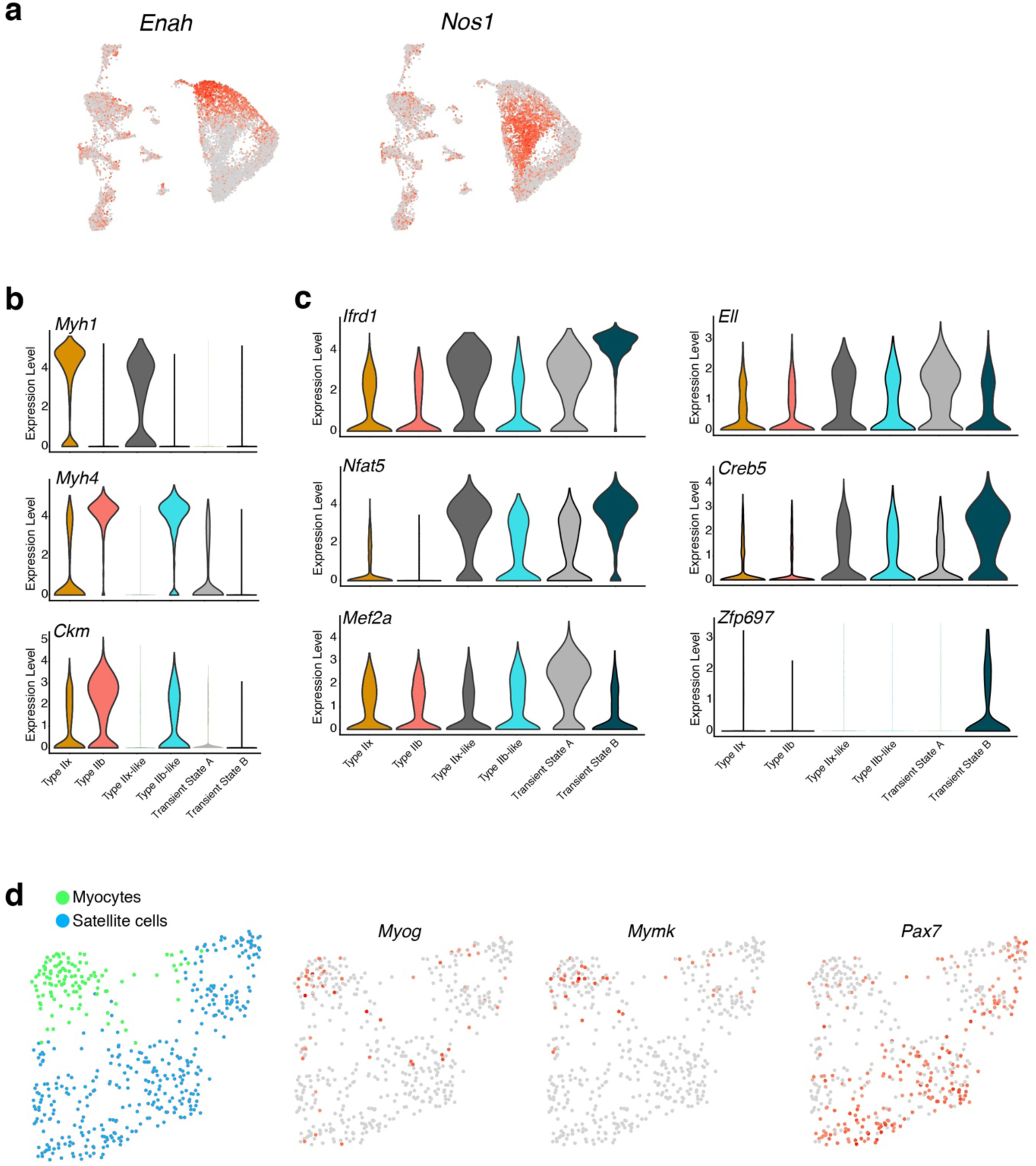
Transient myonuclear states in P21 muscle display non-uniform gene expression. **a**, Feature plots for *Enah* and *Nos1* from the P21 sample. **b**, Violin plots for *Myh1, Myh4*, and *Ckm* show low or minimal expression of these genes in the transient myonuclear populations. **c**, Differential expression of transcription factors in transient myonuclei compared to myonuclei that highly express myosins. **d**, UMAP representation (left panel) after sub-clustering satellite cells and myocytes from the P10 dataset. Panels on right are feature plots for *Myog, Mymk*, and *Pax7*, which confirmed the myocyte population.

**Extended Data Fig. 8.**
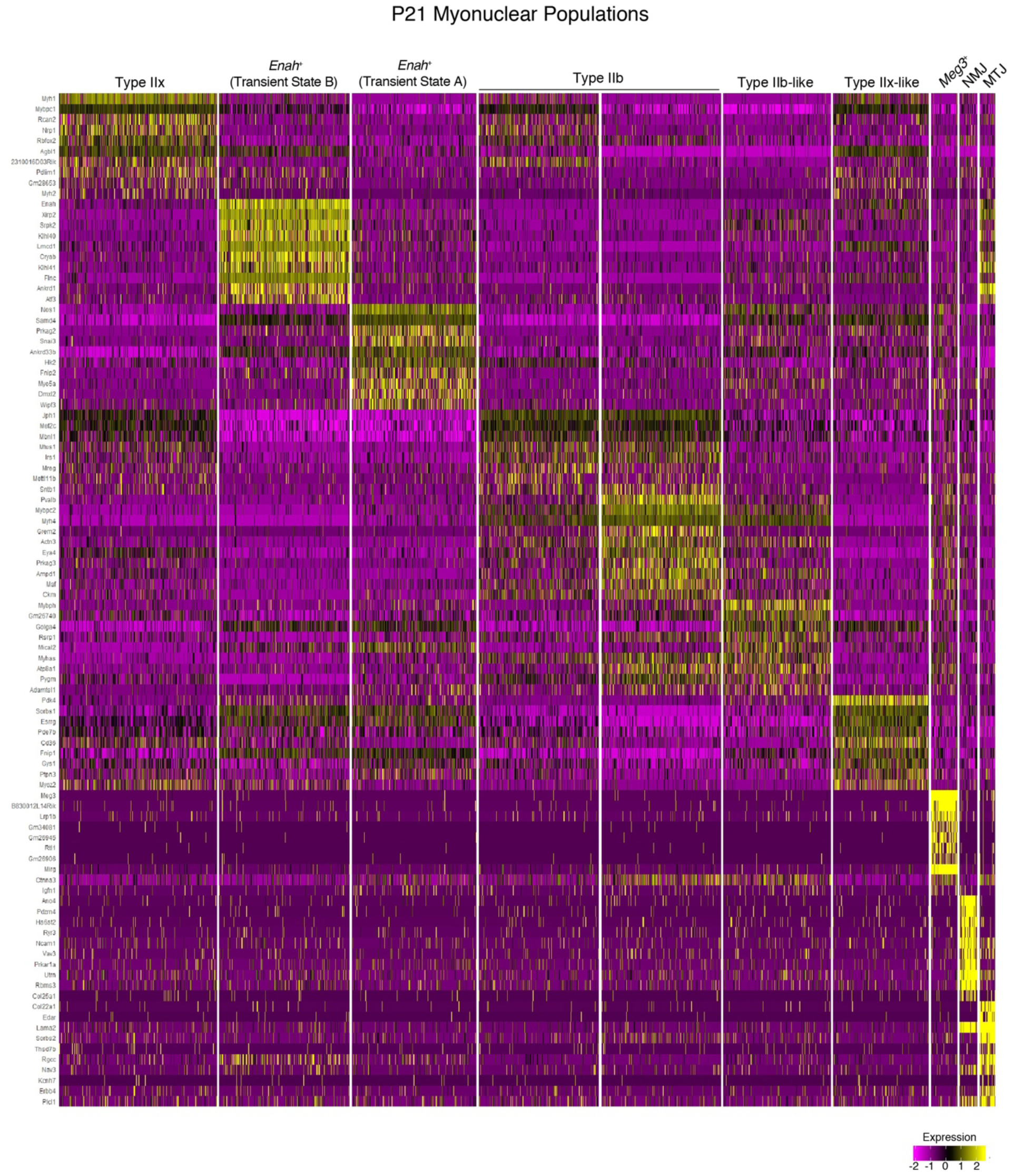
Heterogeneous pattern of skeletal muscle-specific gene signatures in P21 tibialis anterior muscle. Heatmap showing distinct populations of myonuclei in developing muscle, including *Enah*^+^ and *Nos1*^+^ clusters with negligible expression of myosin heavy chains (*Myh4* and *Myh1*) but a distinct enrichment of muscle-specific factors involved in sarcomere formation and myofibrillogenesis.

**Extended Data Fig. 9.**
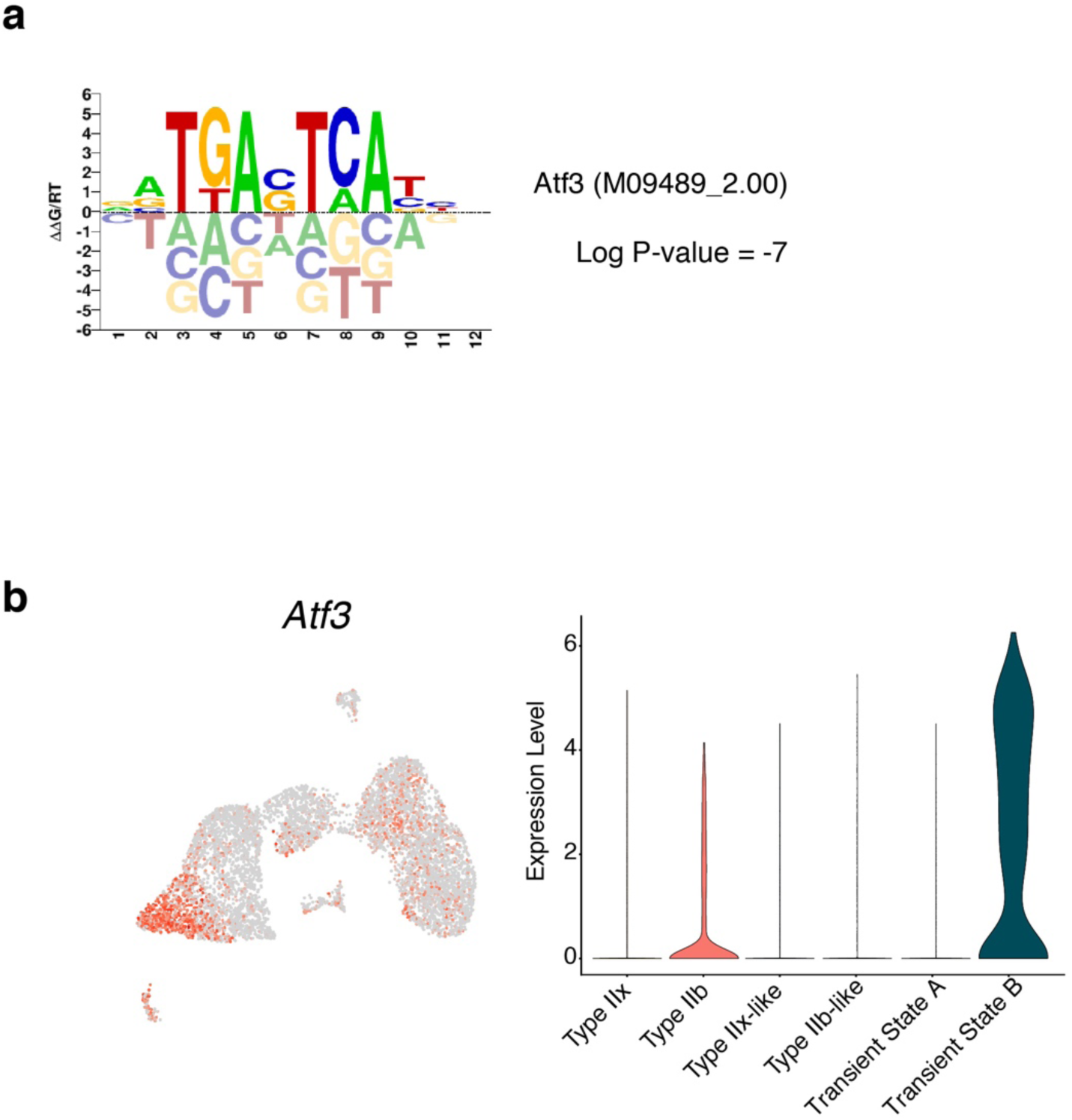
Atf3 as a putative transcriptional regulator of the transient *Enah*^+^ myonuclear program. **a**, Promoter sites of *Enah*^+^ myonuclear marker genes are enriched for Atf3-binding motifs (HOMER motif detection analysis). 30% of genes in the *Enah*^+^ population contain predicted Atf3 binding sites in their promoter region (−1000 – 1000). **b**, Feature plot and violin plot showing that *Atf3* is upregulated at the transcriptional level in *Enah*^+^ transient state B.

**Extended Data Fig. 10.**
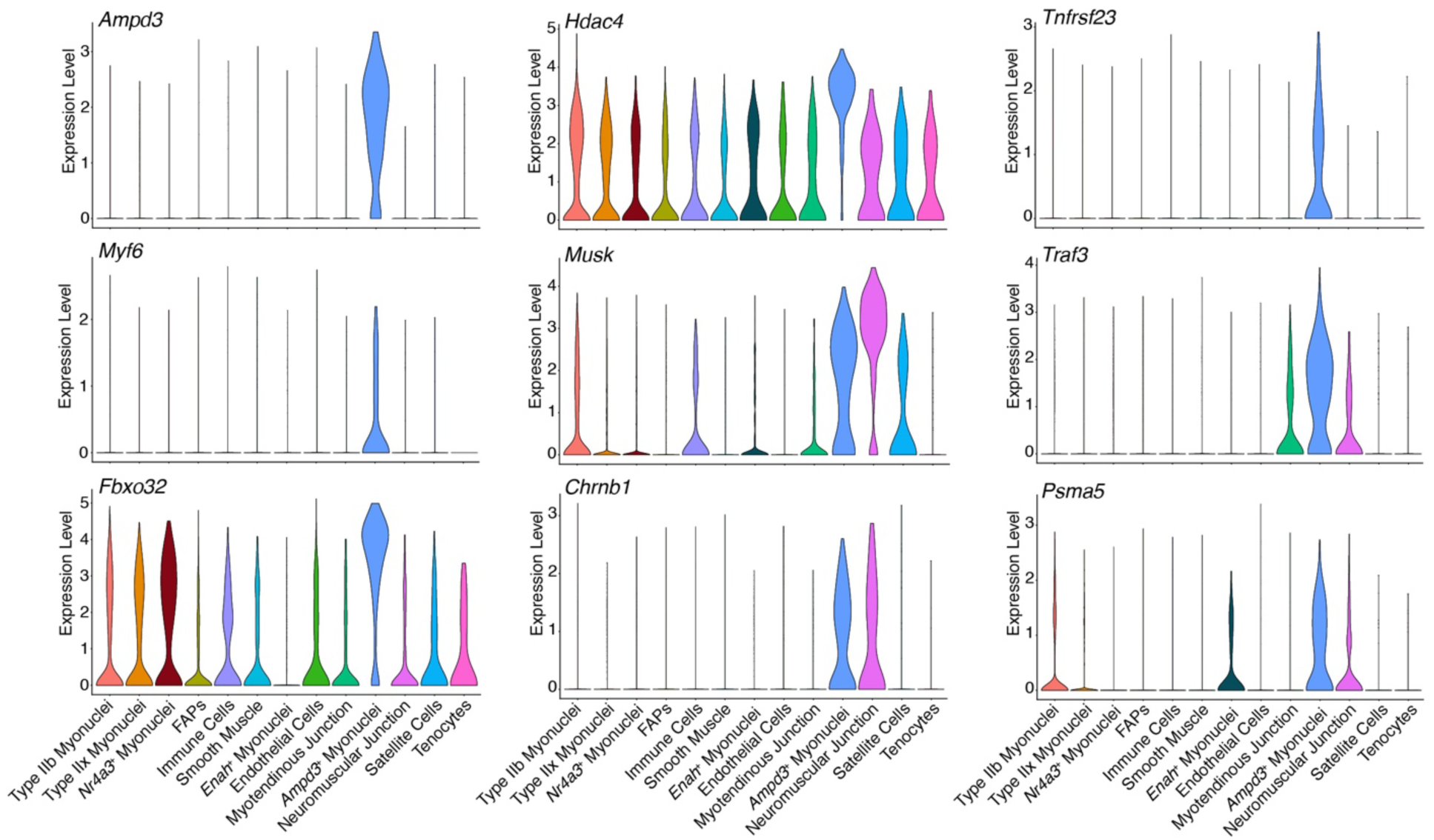
Disruption of gene expression in aged myonuclei. **a**, Violin plots from the nuclei isolated from 30-month old muscle showing *Ampd3*^+^ myonuclei are expressing muscle and NMJ genes, but also are enriched for genes associated with inflammation, cell death, and the proteasome.

**Extended Data Figure 11.**
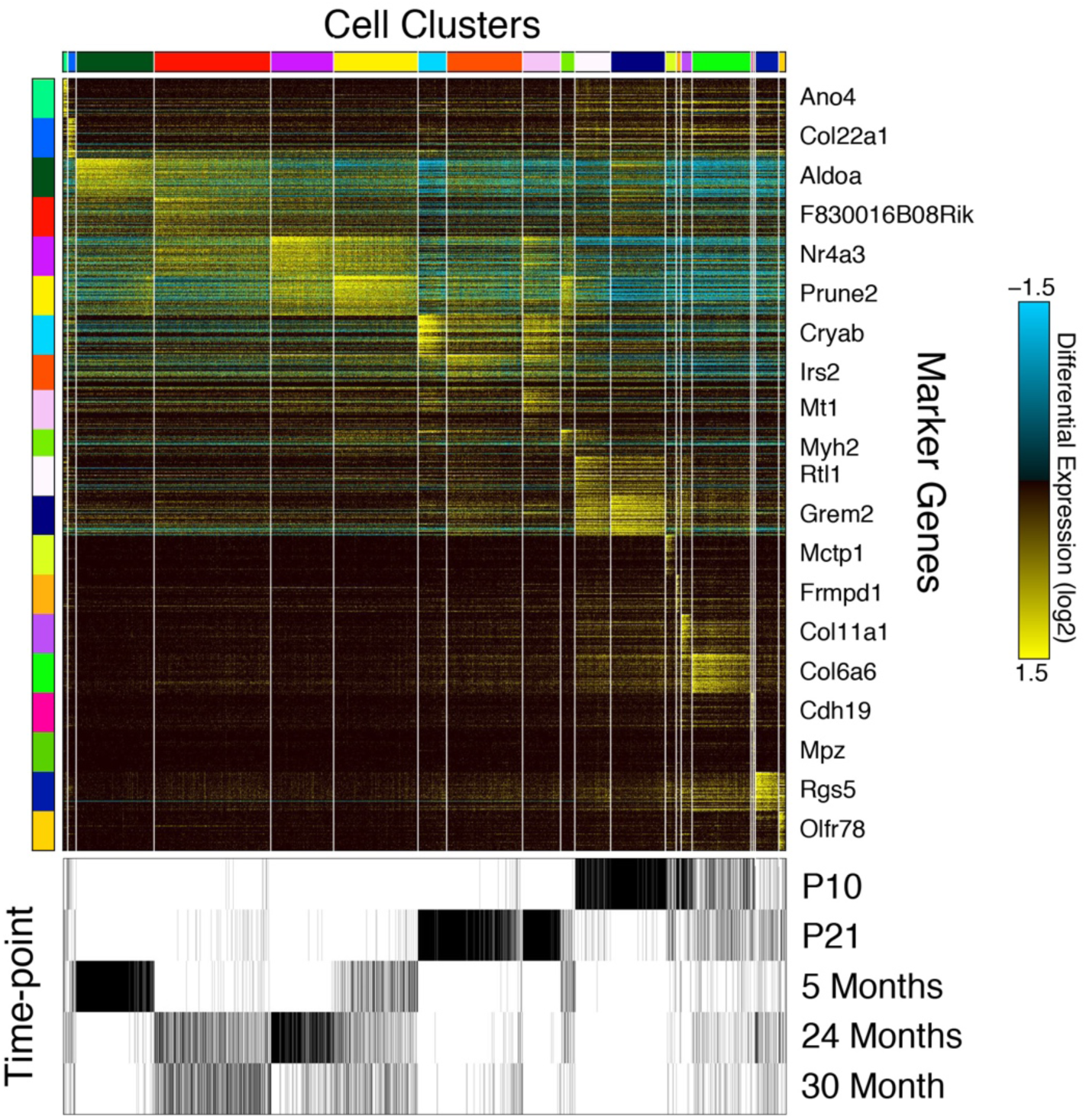
Myonuclei from each time-point intermix to varying degrees based on temporal proximity. Heatmap of gene expression clusters (n=20) identified using the unsupervised ICGS2 workflow of all snRNA-Seq captures. The top 60 marker genes are displayed for each corresponding cell cluster. The time-points associated with each cell are indicated by tickmarks below the heatmap. Enriched gene-sets from diverse single-cell and bulk reference profiles.

## Methods

### Mice and associated experimental procedures

All of the mice used in this study were C57BL/6 wild-type males. For isolation of nuclei for snRNA-seq, tibialis anterior muscles were taken from a single mouse (5 month, 24 month, 30 month) or pooled from 2 mice (P21), or 4 mice (P10) to collect sufficient nuclei for sequencing. Soleus muscles were pooled from 4 mice. Unilateral hindlimb denervation was performed on C57BL/6 wild-type male mice by cutting the left sciatic nerve in the mid-thigh region after the mouse was anesthetized with isoflurane. The gastrocnemius muscle was harvested three days after denervation and the contralateral muscles were used as control. For smFISH experiments, muscles were dissected, embedded in 10% tragacanth/PBS (Sigma), and frozen in 2-methylbutane cooled in liquid nitrogen. 10 μm sections were prepared for all experiments. For RNA isolation, muscles were flash-frozen in liquid nitrogen and stored at -80°C until use. All mouse procedures were approved by Cincinnati Children’s Hospital Medical Center’s Institutional Animal Care and Use Committee (IACUC2017-0053).

### Purification of nuclei from mouse skeletal muscle

Tibialis anterior or soleus muscles were isolated from mice immediately following euthanasia, minced, and placed in homogenization buffer (0.25M sucrose and 1% BSA in Mg^2+^-free, Ca^2+^-free, RNase-free PBS). The minced tissue was further homogenized using an Ultra-Turrax T25. The homogenate was then incubated for 5 min with addition of Triton-X100 (2.5% in RNase-free PBS, added at a 1:6 ratio). Samples were then filtered through a 100 μm strainer, centrifuged (3000 x g for 10 minutes at 4°C) to form a crude pellet, and resuspended in sorting buffer (2% BSA/RNase-free PBS) and filtered again through a 40 μm strainer. Hoechst dye was added to label nuclei as well as 0.2 U/μl Protector RNase inhibitor (Roche). Fluorescently-labeled nuclei were purified via FACS (BD Aria, 70 μm nozzle) and collected in sorting buffer containing RNase inhibitor. The resulting nuclear suspension was pelleted with a light centrifugation step (250 x g for 5 min at 4°C) and the supernatant drawn off to concentrate the nuclei. Aliquots were taken for visualization under a fluorescent microscope to assess nuclear appearance and integrity.

### Construction of libraries and generation of cDNA on the 10X Genomics platform

Nuclei were counted using a hemocytometer and the concentration adjusted if needed to meet the optimal range for loading on the 10X Chromium chip. The nuclei were then loaded into the 10X Chromium system using the Single Cell 3’ Reagent Kit v3 according to the manufacturer’s protocol. Following library construction, libraries were sequenced on the Illumina NovaSeq 6000 system.

### Single-nucleus data analysis

Raw sequencing data of all samples were processed using the cellRanger workflow (version 3.1.0), using a combined intron-exon reference produced as described using the vendor-provided “Generating a Cell Ranger compatible “pre-mRNA” Reference Package” guidelines (https://support.10xgenomics.com/single-cell-gene-expression/software/pipelines/latest/advanced/references). In brief, the “pre-mRNA” reference was derived using the default exon-level GTF file provided by 10x Genomics (http://cf.10xgenomics.com/supp/cell-exp/refdata-cellranger-mm10-3.0.0.tar.gz). Using the below awk command, this exon-level GTF file into “pre-MRNA” GTF containing intron transcript definitions ($ awk ‘BEGIN{FS=“\t”; OFS=“\t”} $3 == “transcript” {$3=“exon”; print}’ genes.gtf > genes.premrna.gtf). Next, the below mkref command was run to produce the final “pre-MRNA” GTF and genome fasta file ($ cellranger mkref --genome=Mmpre --fasta=genome.fa --genes=genes.premrna.gtf). For each dataset, we corrected for ambient background RNA by filtering with the R package SoupX^50^. We used the inferNonExpressedGenes() function to determine which genes had the highest probability of being ambient mRNA, and the strainCells() function in order to transform count matrices. Genes expressed per nucleus was between 1,446 and 2,445, depending on the dataset.

Further data analysis was carried out using the R (version 3.6.1) package Seurat (version 3.1.0)^51^. For individual datasets, outlier nuclei were excluded based on unique feature counts. Nuclei with less than 200 expressed features were excluded, as well as nuclei with greater than 3,200 expressed features (5 month dataset) or 4,000 expressed features (all other datasets). No features that were expressed in 3 or fewer cells were included. Data from filtered nuclei were then normalized logarithmically using the NormalizeData() function. Using the FindVariableFeatures() function, 2,000 features with high variable expression across the nuclei were identified and used in a subsequent Principal Component Analysis (PCA) using the RunPCA() function. For clustering and Uniform Manifold Approximation and Projection (UMAP) visualization, the number of components used was unique to each dataset, based on the strength of PCs visualized by elbow plot and PC heatmaps after PCA was performed. The UMAPs and clusters were generated using the FindNeighbors(), FindClusters(), and RunUMAP() functions (R implementation). Violin and Feature plots were generated using the VlnPlot() and FeaturePlot() from the Seurat package, respectively. For sub-clustering, the function subset() was used to select populations of interest from the data and create a new Seurat object with just those identities, followed by standard Seurat preprocessing. In order to analyze the 5 month dataset with higher dimensionality, the function SCTransform() was used in place of standard preprocessing methods ^15^.

The FindAllMarkers() function was used on each dataset to generate markers for each cluster. Cell-types and nuclear identities were assigned based on a combination of previously reported marker genes along with gene ontology analysis of uniquely expressed genes. Regarding gene ontology, the top 100 marker genes of each cluster were analyzed using ToppGene^52^.

In order to integrate multiple snRNA-seq datasets, the preprocessed datasets to be integrated were run through the FindIntegrationAnchors() function. Resulting integration anchors were used in the IntegrateData() function, creating a new integrated Seurat object. Following this, the standard workflow for visualization and clustering was used. To generate UMAPs of cell identities split or grouped based on their original dataset, the parameters split.by and group.by were set to “orig.ident”. Heatmaps were generated from sub-populations using the DoHeatmap() function. In order to determine differential expression between populations of interest across datasets, the function FindMarkers() was used. In order to identify genes that were consistently changed in aging myonuclei, pairwise comparisons of each aging dataset with the 5-month dataset were performed, and a common aging gene signature was identified. To assess integration of the data, without batch correction, we performed an additional unsupervised analysis using the software ICGS2 (http://altanalyze.org). ICGS2 was run using the default parameters, with cell cycle removal on all combined time-point snRNA-Seq samples.

To identify transcription factor (TF) binding sites that are enriched within the promoter regions of *Nos1*^+^ and *Enah*^+^ genes, we used the HOMER motif enrichment algorithm^53^ and a large library of human position weight matrix (PWM) binding site models obtained from the CisBP database, build 2.0^54^. We used a definition of 1000 bp upstream and downstream of the transcription start site (TSS) to generate consensus predictions.

### Single molecule RNA fluorescent *in situ* hybridization

Single molecule FISH experiments were performed using RNAscope (ACDBio) following the manufacturer’s protocols. We used fresh-frozen cross-sections of quadriceps or gastrocnemius muscles where indicated. Following the completion of the RNAscope fluorescent assay, an immunostaining step was carried out to label myofibers. Sections were blocked using 1% BSA, 1% heat-inactivated goat serum, and 0.025% Tween-20/PBS, followed by incubation for 30 min at RT with anti-laminin (1:100; Sigma L9393). Secondary AlexaFluor antibodies (1:200) (Invitrogen) were then applied at room temperature for 1 hour. For visualization of neuromuscular junctions, an α-Bungarotoxin AlexaFluor conjugate (1:100; Thermo Fisher) was added with the secondary antibody. Sections were mounted using VectaShield with DAPI (Vector Laboratories). Slides were imaged using a Nikon A1R confocal system.

### Cell culture

C2C12 cells were acquired from American Type Culture Collection. Cells were grown in DMEM (Gibco) containing 10% heat-inactivated bovine growth serum (BGS) and supplemented with penicillin-streptomycin (1%), and incubated at 37°C with 5% CO2. C2C12 cells were differentiated by switching medium to DMEM with 2% heat-inactivated horse serum with antibiotics. In order to induce formation of aneural AChR clusters, 35mm plates were pre-coated with 10 μg/ml natural mouse laminin protein (ThermoFisher 23017015) suspended in L-15 medium (Thermo Fisher) with 0.2% NaHCO3, as previously described^47^. Cells were transfected with siRNAs on day 2 of differentiation using Lipofectamine 2000 (Thermo Fisher) according to the manufacturer’s protocol. All siRNAs were purchased from Santa Cruz.

On day 5 of differentiation, cells were rinsed with PBS and processed for RNA isolation or immunocytochemistry. For the latter, cells were fixed in 4% paraformaldehyde for 20 minutes at room temperature. Cells were then permeabilized with 0.2% Triton X-100/PBS for 20 minutes at room temperature, followed by blocking in 3% BSA/PBS for 30 minutes. We then incubated with phalloidin (1:100, Thermo Fisher) and α-Bungarotoxin (1:200, Thermo Fisher) AlexaFluor conjugates for 1 hour at room temperature. Cells were then rinsed with PBS and nuclei were stained with Hoechst 33342 solution (Thermo Fisher). Imaging was performed using an upright Nikon FN 1 microscope on a Nikon A1R confocal.

### Quantitative real-time PCR

RNA was isolated from homogenized muscle tissue and plated cells using Trizol reagent (Invitrogen) according to standard protocols. cDNA synthesis was carried out using the MultiScribe™ kit (Applied Biosystems) and qPCR reactions were performed on a Bio-Rad CFX96™ Real-Time System using PowerUp™ SYBR Green Master Mix (Applied Biosystems). Primers used for qPCR are listed below.

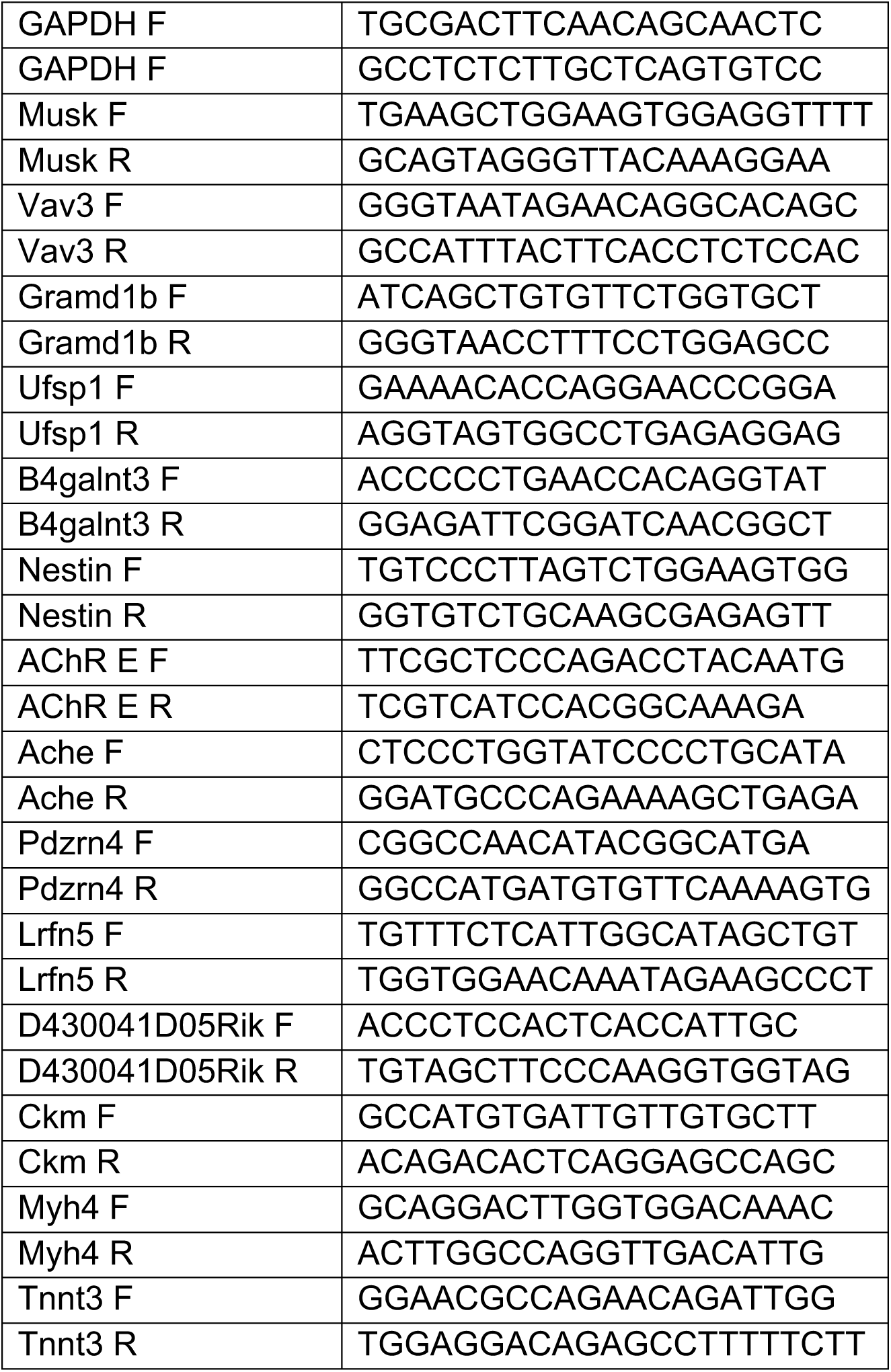

### Statistics

GraphPad Prism 8 software was used for statistical analysis, excluding single nucleus RNA-sequencing data. Unpaired t-tests were used to determine statistical significance, which was set at p values <0.05. For qPCR analysis, ΔCt values were used to assess significance, and ΔΔCt values were graphed to show relative expression compared to control.

### Data availability

snRNA-Seq data processed data, cell populations and quality control information has been deposited in Synapse (https://www.synapse.org/#!Synapse:syn21676145). Reviewers may access the Synapse data using the Username: tempreviewer1 and Password: l6GawO3N. Raw sequencing data are being deposited in GEO and will be publicly available after publication.

### Code availability

Data analysis was limited to standard pipelines in both Seurat and SoupX R packages, as described in the Methods. Scripts will be made available on GitHub after publication.

## Acknowledgements

We thank the following entities at Cincinnati Children’s Hospital Medical Center: Kelly Rangel and Shawn Smith from the Gene Expression Core, David Fletcher at the Sequencing Core, Steven Potter, and Hee-Woong Lim. The main funding source for this work was from the Research Innovation and Pilot Funding Program of the Cincinnati Children’s Hospital Research Foundation to D.P.M. and N.S. This work was also supported by grants to D.P.M. from the National Institutes of Health (R01AR068286, R01AG059605) and Pew Charitable Trusts. A Cincinnati Children’s Hospital Endowed Scholar Award supported D.P.M. and M.T.W. M.T.W. laboratory was also supported by grants from the National Institutes of Health (R01NS099068, R01AR073228, R01GM055479). C.O.S. was supported by an undergraduate fellowship from the American Heart Association (18UFEL33930019). M.J.P. was supported by a training grant from the National Institutes of Health (NHLBI T32HL007752).

## Author contributions

M.J.P. and D.P.M. designed experiments. M.J.P., C.O.S., C.Su., X.C., and K.C. performed experiments. M.T.W. and N.S. provided bioinformatic guidance. All authors analyzed the data and contributed to interpretations. M.J.P. and D.P.M. wrote the manuscript with assistance from all authors.

